# Your Blood is Out for Delivery: Considerations of Shipping Time and Temperature on Degradation of RNA from Stabilized Whole Blood

**DOI:** 10.1101/2024.08.24.609519

**Authors:** Filip Stefanovic, Lauren G. Brown, James MacDonald, Theo Bammler, Darawan Rinchai, Serena Nguyen, Yuting Zeng, Victoria Shinkawa, Karen Adams, Damien Chausabel, Erwin Berthier, Amanda J. Haack, Ashleigh B. Theberge

**Affiliations:** Department of Chemistry, University of Washington, Seattle, Washington 98195, United States; Department of Environmental and Occupational Health Sciences, University of Washington, Seattle, Washington 98195, United States; Department of Infectious Diseases, St Jude’s Children Research Hospital, TN, Memphis 38105, United States; Institute of Translational Health Sciences, School of Medicine, University of Washington, Seattle, Washington 98195, United States; Computer Sciences Department, The Jackson Laboratory, Farmington, CT, 06032, United States; School of Medicine, University of Washington, Seattle, Washington 98195, United States; Department of Urology, School of Medicine, University of Washington, Seattle, Washington 98195, United States

## Abstract

Remote research studies are an invaluable tool for reaching populations in geographical regions with limited access to large medical centers or universities. To expand the remote study toolkit, we have previously developed homeRNA, which allows for at-home self-collection and stabilization of blood and demonstrated the feasibility of using homeRNA in high temperature climates. Here, we expand upon this work through a systematic study exploring the effects of high temperature on RNA integrity through in-lab and field experiments. Compared to the frozen controls (overall mean RIN of 8.2, *n* = 8), samples kept at 37°C for 2, 4, and 8 days had mean RINs of 7.6, 5.9, and 5.2 (*n* = 3), respectively, indicating that typical shipping conditions (∼2 days) yield samples suitable for downstream RNA sequencing. Shorter time intervals (6 hours) resulted in minimal RNA degradation (median RIN of 6.4, *n* = 3) even at higher temperatures (50°C) compared to the frozen control (mean RIN of 7.8, *n* = 3). Additionally, we shipped homeRNA-stabilized blood from a single donor to 14 different states and back during the summer with continuous temperature probes (7.1 median RIN, *n* = 42). Samples from all locations were analyzed with 3’ mRNA-seq to assess differences in gene counts, with the transcriptomic data suggesting that there was no preferential degradation of transcripts as a result of different shipping times, temperatures, and regions. Overall, our data support that homeRNA can be used in elevated temperature conditions, enabling decentralized sample collection for telemedicine, global health, and clinical research.

## INTRODUCTION

Remote self-sampling studies have become more prevalent in the wake of the COVID-19 pandemic, however, their utility goes far beyond the convenience of at-home sampling in a pandemic setting.^1–5^ Fully remote studies circumvent many logistical barriers that preclude underserved and rural populations from participating in clinical research. These barriers include access to clinics or study sites that can perform blood draws, need for trained phlebotomists, suitable transportation, and sufficient time outside of typical work schedules. Additionally, remote sampling is compatible with collection of time-sensitive and longitudinal samples, making it an invaluable tool for human health research.^6–11^

Devices for remote blood sampling (such as lancet-based devices from Tasso (Seattle, WA), YourBio Health (Medford, MA), etc.) offer a user-friendly way to self-collect blood samples. To enable analysis of many blood analytes, remote sample stabilization is necessary, particularly for whole blood RNA. Without stabilization, RNAses in whole blood can trigger intracellular RNA transcript degradation pathways *ex vivo* that can alter gene expression levels.^12–15^ To address this aspect of RNA degradation, our lab has developed the homeRNA kit which consists of a commercially available Tasso-SST upper-arm blood collection device (which collects up to 0.5 mL of blood) and a custom-engineered stabilizer tube containing RNA*later*.^9^ RNA*later* is a commercially available RNA stabilization agent commonly used with biological tissue samples that inhibits RNase activity and stabilizes cellular RNA.^16–19^ In previous homeRNA studies, we have demonstrated that RNA*later* effectively stabilizes self-collected blood and yields RNA of sufficient quality and quantity for downstream transcriptome analyses.^9–11,20^ In these studies, samples experience variable shipping times and often are not stored in the freezer until >48 h after collection. In ongoing work, we use homeRNA to study the inflammatory response to wildfire smoke exposure, which includes sampling during hot summer months across Western and Central U.S.,^21^ and acute immune response to COVID-19 infection with nationwide sampling during all seasons.^10,11^ Since the US has vastly different climates depending on the time of year and location, it is important to consider how the shipping process may affect the integrity of RNA in remotely collected and stabilized blood samples. For example, our wildfire smoke exposure study took place over a 10-month period (before, during, and after wildfire smoke exposure) and many samples were collected during the summer months. Further, we have previously conducted a preliminary examination of homeRNA in the Western and Central U.S. and Qatar with the goal of validating its utility in high temperature settings.^20^

While the majority of samples collected in our previous and ongoing studies have sufficient RNA quality and yield for transcriptomic analysis, it is important to further investigate if transcripts are preferentially degraded and could bias the interpretation of these results. A potential concern based on past remote study experiences is that rural communities experience longer shipping times which could adversely affect RNA quality. Similarly, there is a concern that remote studies investigating immune response in tropical climates or during summer months could suffer from heat-induced RNA degradation.

The existing body of research has primarily investigated the effects of cold and ambient temperatures on RNA integrity prior to isolation from whole blood samples, as well as stability of RNA following extraction.^22–25^ However, few studies to date have systematically investigated exposure of blood to high temperatures (>37°C) and its effect on the resulting isolated RNA quality; one study from Sarathkumara et al. investigated the effect of exposing Tempus- and PAXgene-stabilized blood samples up to 40°C and found a decrease in RIN with prolonged exposure (up to 10 days).^26^ Additionally, Heneghan et al. observed a 5-10 fold increase in the rate of RNA transcript degradation from dried blood spots stored at 37°C, suggesting that higher temperatures may compromise the integrity of the transcriptome in dried blood spots.^27^ Drawing on our past experiences with homeRNA and remote studies, we designed a set of temperature-controlled experiments on RNA*later*-stabilized whole blood samples exposed to temperatures up to 50°C for up to 8 days. Additionally, we conducted a real-world shipping experiment and sequenced a subset of these samples using 3’ mRNA sequencing (3’ mRNA-seq) to better understand how variable exposure to different temperatures and shipping times can affect the quality of the transcriptomic data. This work serves as a roadmap for developing future remote transcriptomic studies that take place in elevated temperatures (e.g., tropical climates, summer months) and lays the groundwork for understanding the role temperature degradation plays in the interpretation of these data.

## EXPERIMENTAL

### Investigating RNA Integrity of RNA*later* Stabilized Blood Samples in Temperature Controlled Experiments

#### Exposure to Longer Term (2, 4, and 8 days) High Temperatures (>37°C)

A venous blood draw was performed and promptly stabilized with RNA*later* at the manufacturer’s recommended ratio (2.6 mL RNA*later* per 1 mL blood). For the first biological replicate, blood was acquired via venipuncture from Bloodworks and stabilized within 1-2 hours. For the second and third replicate, blood was collected via venipuncture in lab under study number STUDY00014133. After stabilization with RNA*later*, the stabilized blood was aliquoted in 1.8 mL samples to replicate the maximum volume of homeRNA (0.5 mL blood and 1.3 mL RNA*later*). After stabilization, each sample was stored according to the following conditions: (1) immediately incubated at set temperatures (25°C, 37°C, 40°C, 45°C, and 50°C) for 2, 4, and 8 days, (2) kept at 4°C for 24 hours then incubated at set temperatures (25°C, 37°C, 40°C, 45°C, 50°C) for 2, 4, and 8 days, and (3) kept at 25°C for 24 hours then incubated at set temperatures (25°C, 37°C, 40°C, 45°C, 50°C) for 2, 4, and 8 days. Each condition had a corresponding control: (1) frozen immediately at -20°C, (2) incubated at 4°C for 24 hours then frozen at - 20°C, (3) incubated at 25°C for 24 h then frozen at -20°C. One sample at each temperature and condition was removed after 2, 4, and 8 days such that there was a single replicate for each temperature and condition on each day of removal. At these timepoints, the blood was moved from the incubators to the -20°C freezer until ready for RNA isolation. For each condition, RNA from the stabilized blood samples was extracted based on timepoints, resulting in three total extraction batches based on day removed after high temperature exposure (day 2, 4, and 8) with six samples in each batch (condition control, 25°C, 37°C, 40°C, 45°C, and 50°C). The experiment was repeated two more times for a total of three replicates from two donors (replicate 2 and 3 from the same donor, with the blood drawn on separate days).

#### Exposure to Shorter Term (<2 days) High Temperatures (>37°C)

A venous blood draw was performed in lab under study number STUDY00014133 and promptly stabilized with RNA*later* at the manufacturer’s recommended ratio (2.6 mL RNA*later* per 1 mL blood). The stabilized blood was then aliquoted to 1.8 mL samples. One sample was immediately frozen at -20°C as the control. The rest of the samples were then treated according to the following conditions: (1) one sample kept at room temperature (25°C) overnight (16 h), then frozen (−20°C), (2) five samples kept at room temperature (25°C) overnight (16 h), moved to incubators of corresponding temperatures (25°C, 37°C, 40°C, 45°C, 50°C) for 6 h, then frozen (−20°C), (3) five samples kept at room temperature (25°C) overnight (16 h), moved to incubators of corresponding temperatures (25°C, 37°C, 40°C, 45°C, 50°C) for 6 h, moved back to room temp (25°C) for 24 h, then frozen (−20°C). This resulted in a total of one replicate for each temperature and condition combination. The stabilized blood samples were stored at -20°C until ready for RNA isolation and assessment. All 12 samples from each replicate were extracted in one batch. The experiment was repeated two more times for a total of three replicates from three separate blood draws from one donor.

### Investigating RNA Integrity of RNA*later* Stabilized Blood Samples in Real World Setting with homeRNA

#### Shipping of RNAlater-Stabilized Blood Across United States with Continuous Temperature Monitoring

A venous blood draw was performed in the lab under study number STUDY00014133. The blood sample was aliquoted into 48 Tasso-SST tubes at 0.5 mL each. The blood was then stabilized by attaching the Tasso-SST tubes to a custom-designed stabilizer tube with 1.3 mL RNA*later* and shaken to mimic blood stabilization in homeRNA kits. The remaining blood was stabilized in bulk (2.6 mL RNA later per 1 mL blood), aliquoted into 1.8 mL samples, and immediately frozen at -20°C as controls. The 48 homeRNA-stabilized samples were then placed in 50 mL conical tubes fitted with custom adapters as described in Haack et al. 2021.^9^ The 50 mL tubes were then adhered together using tape in sets of three such that three technical replicates would be sent to each shipped location. Each set had an RC-5+ continuous temperature monitor (Elitech) directly attached with lab tape (see Figure S1A), with temperatures being recorded every 2 minutes. These constructs were then placed in a biohazard bag and into the box used for homeRNA kits (see Haack et al., 2021) prior to being packaged into a UPS LabPak. The full setup is shown in Figure S1. The complete packages were shipped to volunteers in 14 different states across the United States via UPS. The states included: Washington (WA), New Mexico (NM), North Carolina (NC), Minnesota (MN), Maine (ME), Massachusetts (MA), Kansas (KS), Illinois (IL), Georgia (GA), Colorado (CO), California (CA), Arizona (AZ), Hawaii (HI), and Nebraska (NE). One additional package was kept in a 25°C incubator (In-Lab/Lab). Upon delivery, the volunteers were instructed to keep the packages inside overnight at room temperature and adhere the return labels to the package. For pickup, most volunteers left the package outside on the front porch, two volunteers left the package indoors in a mailroom, and one volunteer directly handed the package to the UPS courier. Most samples were picked up within 1 – 2 days from the volunteer and shipped back to the lab. Shipping return times ranged from overnight to 4 days after pickup. Upon return to the lab, the packages were immediately placed into a -20°C freezer. Once all the packages were returned, the samples that were kept in the lab in a 25°C incubator were also frozen at -20°C. All samples were stored at -20°C until ready for RNA isolation. RNA was extracted from all shipped homeRNA-stabilized blood samples in three extraction batches, with one technical replicate from each shipping location and in-lab baseline samples (immediately frozen and constant 25°C sample), resulting in 16 samples extracted in each batch.

### RNA Sequencing and Analysis

#### 3’ mRNA-seq of Real-World Shipping Experiment

After isolation the samples were tested for RIN scores on the Agilent 2100 Bioanalyzer (see Supplemental Information for methods and Figure S8 for resulting electropherograms). A 100 ng of total RNA from 20 selected samples were sequenced by the Lexogen facility (Vienna, Austria) using 3’ mRNA-sequencing technology (Lexogen QuantSeq 3’ mRNA-seq V2 Library Prep Kit FWD with UDI) on an Illumina NextSeq 2000 instrument according to the manufacturers’ standard instructions. The 16 samples from the third batch were selected for sequencing as the third batch had the greatest proportion of the median RIN values amongst the three replicates from each location. Additionally, all three replicates from the baseline sample (frozen immediately after stabilization) and all three replicates from the location with the lowest RIN (NE, mean RIN of 5.5) were sent for sequencing. In total, 20 samples (16 from batch 3, 2 additional from NE, 2 additional from In-Lab) were sent for 3’ mRNA sequencing. Prior to library preparation, all samples were treated with Ambion DNase I (Invitrogen) for 10 minutes at 37°C followed by heat inactivation with EDTA at 75C for 10 minutes. Libraries were prepared using 17 ng of RNA as input. Libraries were amplified for 18 cycles and were quality controlled on a Fragment Analyzer device using the DNF-474 HS NGS Fragment kit (1-6000 bp) (Agilent). The samples were sequenced to a read depth of 5 million reads.

#### Module Enrichment Score Calculation

Enrichment scores were calculated using the gene set variation analysis (GSVA) method for each of the 382 transcriptional modules from the BloodGen3 repertoire across all samples.^28,29^

#### Principal Component Analysis

Principal component analysis (PCA) was performed on the module enrichment scores to visualize the variance among samples, labeled by their condition (shipping location, in-lab sample, or control).

#### Heatmap Visualization

Module enrichment scores were represented in a heatmap format, with rows corresponding to different modules associated with biological processes such as interferon response and inflammation, and columns representing individual samples. The color gradient of the heatmap indicates the level of enrichment, ranging from decreased (blue) to increased (red).

#### Dot Plot Visualization

For a more detailed analysis, we focused on specific module groups (A28, A35, and A37) and plotted their enrichment scores using dot plots. The x-axis represents individual samples, while the y-axis denotes the corresponding enrichment score for each sample within the selected modules.

#### Interferon Response Kinetics Analysis

We used data from the COVAX study (Rinchai et al 2022), to illustrate the kinetics of six interferon response modules (M8.3, M10.1, M13.17, M15.64, M15.86, and M15.127) at different timepoints before and after administration of COVID-19 mRNA vaccine.^30^ Enrichment scores for these modules were plotted for each sample across the various timepoints.

#### Transcriptomic Analysis of Individual Gene Expression Counts

Summarized counts/genes were imported into R and analyzed using the limma-voom pipeline, which uses conventional weighted linear regression to model the data.^31,32^ The matrix of gene counts was filtered to remove those genes with consistently low expression, and then converted to log counts/million counts, using estimated library sizes calculated using the trimmed mean of M-values (TMM) method.^33^ The logCPM values have a dependence between the mean and variance which is estimated as part of the limma-voom pipeline and used to estimate observation-level weights. These weights are used as part of the weighted line regression to adjust for heteroskedasticity. We fit a model that includes time >30°C for each sample, the RIN (to adjust for RNA quality), and five surrogate variables computed using the Bioconductor sva package. The surrogate variables are intended to adjust for unobserved variability, and have been shown to increase power to detect differences.^34^ We then tested for any apparent changes in gene expression that correlate with exposure to high temperatures.

## RESULTS AND DISCUSSION

### Investigating RNA Integrity of RNAlater Stabilized Blood Samples in Temperature Controlled Experiments

To validate the utility of the homeRNA kit for preservation of RNA, we conducted a set of experiments to systematically probe the effects of storage conditions (time, temperature, repeat exposure). We chose times and temperatures to replicate scenarios similar to what samples would experience in a remote study conducted with the homeRNA platform.^9–11,20^ In a typical homeRNA study, participants are given instructions for self-collection and blood stabilization. Participants then package their stabilized samples for next- or same-day courier pickup. Many study participants opt to leave the samples on their front porch for a pickup window of 2 – 4 hours. Based on the location and time of year, samples may experience multiple hours outdoors in hot temperatures (>37°C). After pickup, packages typically take 1 – 2 days to be returned to the lab, depending on the participant’s location; but in rare cases it has taken up to 2 weeks for a package to be returned.^9^ Additionally, while in transit, samples may experience fluctuations in temperatures (e.g., prior to pickup, during transit, or in storage facilities) that could affect the stability of the RNA and cause differential degradation of RNA transcripts (Figure S2). In this work, we aim to elucidate (1) if exposure to longer shipping times and higher temperatures results in increased degradation of RNA and (2) if the variable conditions in the shipping process result in preferential degradation of specific transcripts.

To model shipping times and temperatures that homeRNA samples may experience, we exposed RNA*later*-stabilized venous blood samples to various temperatures (25°C, 37°C, 40°C, 45°C, or 50°C) and varying lengths of time (6 h, ∼1 day, 2 days, 4 days, or 8 days) in the lab. We place emphasis on the effects of higher temperature conditions (>37°C) since some of our ongoing homeRNA studies take place in hot climates or during the summer months. Remote studies can also take place in winter months or colder climates and may experience freezing with subsequent thawing prior to processing. Although some literature reports adverse effects of freeze-thaw cycles on RNA integrity from whole blood samples, we have found that RNA*later*-stabilized whole blood samples are minimally affected by a freeze-thaw cycle (Figure S3). ^22,35,36^

### Longer Term (2, 4, and 8 day) Exposure to High Temperatures (>37°C)

Our first experiment was set up to understand the effect of “longer-term” exposure to various temperatures and storage times (2, 4, or 8 days) of the stabilized blood samples and the resulting effects on RNA integrity. These time points were chosen to recapitulate potential scenarios homeRNA samples may experience. In our previous remote studies, most samples were returned within 1 – 2 days of collection, with some samples taking 4 days or more due to logistical complications or weather events (wildfire, storms, etc.).^9–11^ To capture a “worst-case” scenario, we included a longer time point (8 days). While we have opted for overnight shipping with private companies (UPS, FedEx) to minimize time samples spend in transit, we recognize that these are not available to all researchers who may want to use the homeRNA platform. As such, the longer time points (8 days) are valuable for an array of scenarios where samples spend extended periods in transit such as overseas shipping, or shipping from rural areas. Additionally, we chose to hold the samples at steady temperatures (from 25°C to 50°C, Figure 1) to understand temperature-dependent degradation of RNA. For this experiment, we chose three conditions to represent three different scenarios: (condition 1) the sample is left out on the porch for pickup immediately after collection, (condition 2) the sample is kept in a fridge prior to being left on the porch for pickup, and (condition 3) the sample is kept indoors (not refrigerated) overnight prior to being left outside for pickup. All samples were stored at -20°C until being extracted. We note here that RNA*later*-stabilized samples are known to be stable indefinitely at -20°C in the product’s official documentation and that samples are often temporarily stored in a freezer at this temperature upon delivery.^37^

**Figure 1.**
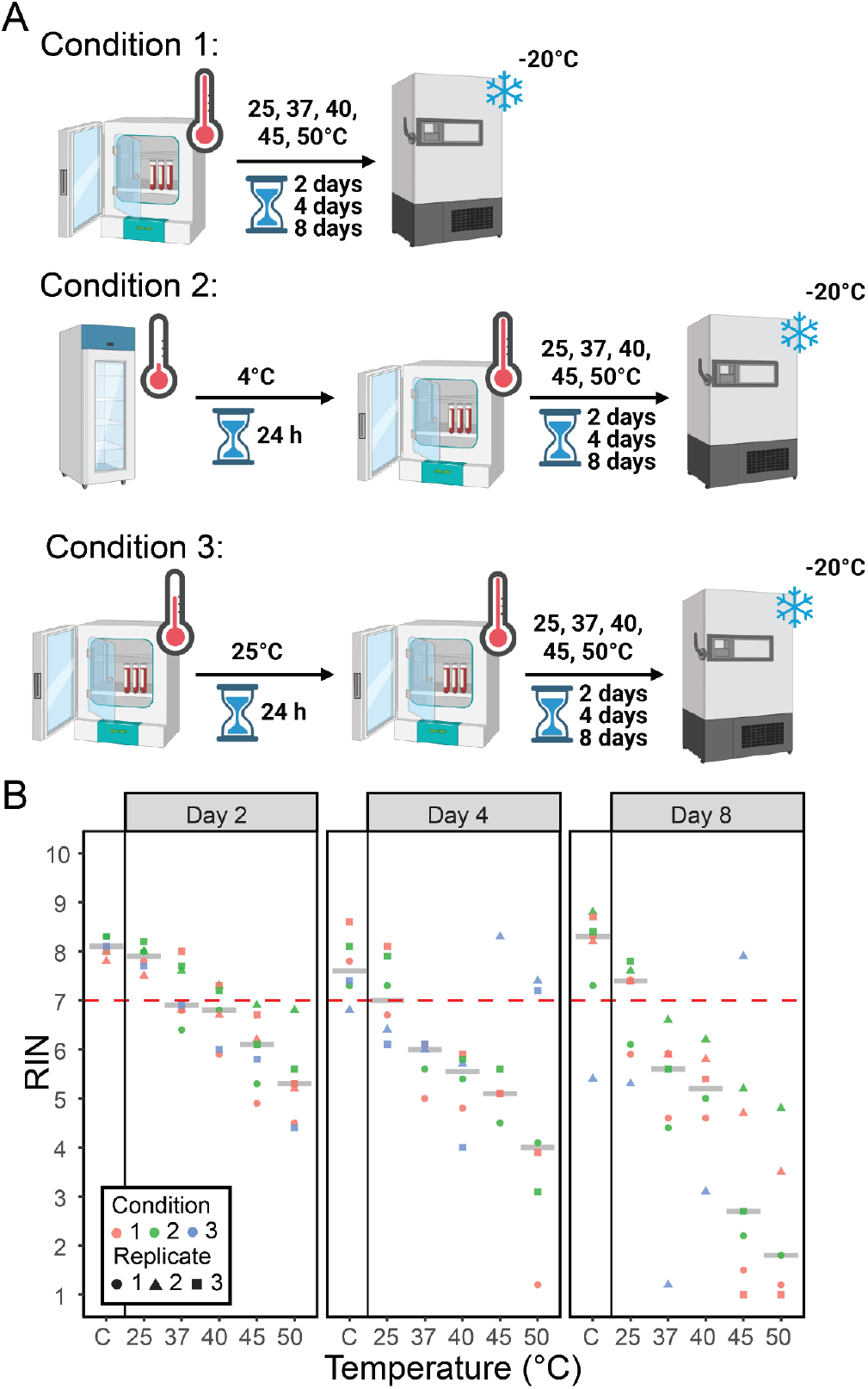
Exposure of RNA*later*-stabilized whole blood samples to longer term (2, 4, and 8 day) high temperatures (>37°C). (A) Outline of experimental conditions. Condition 1, samples were immediately incubated at 25°C, 37°C, 40°C, 45°C, and 50°C for 2, 4, and 8 days. One sample was frozen at -20°C immediately after stabilizing with RNA*later*; Condition 2, samples were incubated at 4°C for 24 hours then exposed to the same set of temperatures and times as condition 1. One sample was frozen at -20°C after incubation for 24 hours at 4°C; and condition 3, samples were incubated at 25°C for 24 hours prior to exposure to the same set of temperatures and times as condition 1. One sample was frozen at -20°C after incubation for 24 hours at 25°C. After each timepoint, one sample from every temperature was immediately frozen and kept at -20°C until ready for RNA extraction. Created with BioRender.com. (B) RNA quality of each extracted sample from each condition. For each condition, samples were extracted based on timepoints, resulting in three total extraction batches (day 2, 4, and 8) with 6 samples in each batch (condition control, 25°C, 37°C, 40°C, 45°C, and 50°C). Each condition was performed in triplicate. The gray crossbars signify the mean RIN at each time and temperature across all conditions and replicates.

RNA Integrity Number (RIN) is a common metric used to measure the quality of RNA samples.^38,39^ There are some discrepancies in the literature about a cutoff RIN value that is suitable for sequencing, with values of 7 or 8 often cited.^14,40–42^ However, sequencing technologies are evolving at a rapid pace, and the interpretation of RIN is far more nuanced than using a single number to evaluate the usability of a given RNA sample. Newer sequencing technologies such as 3’ mRNA-seq and post-sequencing computational processing methods make possible the sequencing of samples with RINs as low as 3.^14,41,43–45^ Our data show that increasing temperatures results in greater RNA degradation (lower RIN values) in RNA*later*-stabilized blood samples as compared to samples immediately frozen after stabilization (Figure 1). Longer exposures resulted in additional degradation; for example, the mean RIN for the 50°C condition changed from 5.5 to 4.0 to 1.8 for the samples at 2, 4, and 8 days, respectively. When incubated for two days at any temperature, all samples had RINs that were suitable for downstream transcriptomic analysis (from 7.9 at 25°C to 5.5 at 50°C). Although notable degradation was observed for samples that were kept at 45°C and 50°C for 8 days, it is also important to note that samples in a remote study will be subject to variable temperatures and not a constant exposure to the high temperatures over several days. For example, it is highly unlikely that a sample would be exposed to 8 days at 50°C, the highest and longest condition that we tested. Moreover, samples that were kept at 25°C had a mean RIN of at least 7 across all time points, suggesting that RNA*later*-stabilized samples at lower temperatures undergo limited degradation even on longer time scales (up to 8 days). Further, samples in condition 2 (refrigerated prior to shipment) gave similar RIN values as samples in condition 3 (kept at room temperature overnight). Taken together, these results suggest that the additional refrigeration step prior to sample shipping is not strictly necessary.

### Shorter Term (<2 days) Exposure to High Temperatures (>37°C)

Beyond investigating longer times samples may experience at high temperatures (>37°C) during shipping, we were also interested in shorter durations (within hours) of high temperature exposure. In homeRNA studies, participants will often opt to have the sample picked up by a courier service by leaving the sample on their front porch for pickup. This means that a study participant may leave their sample outside before leaving for work at 8 AM and the package may not get picked up until later in the day. Depending on the time of year and location, the sample may be exposed to very high temperatures for several hours. In the shipping experiment (see section below), we left the samples in a UPS box at the University of Washington prior to pickup. The temperature monitors in these packages showed a peak during this time, with an average maximum temperature of 38.6°C (over 14 temperature probes) and reaching up to 41.9°C (one temperature probe). Here the samples were not exposed to warm temperatures for a full 2 – 8 days but rather only for 6 – 8 hours. Given these two scenarios where samples have brief exposures to peak temperatures, we wanted to investigate the effect of high temperature on a shorter time scale.

To better mimic the conditions homeRNA samples may experience in the context of remote studies, samples were exposed to conditions replicating different scenarios prior to shipment. All experimental samples were kept at room temperature (25°C) overnight (16 hours) to emulate a study participant collecting their sample and leaving it in their home until the next day’s pickup. One stabilized sample was frozen immediately as the control. After the initial overnight room temperature incubation, the experimental samples were either frozen or exposed to the same set of temperatures established in our previous longer-term experiment (25°C, 37°C, 40°C, 45°C, and 50°C) for 6 hours. The samples that were frozen after incubation at room temperature overnight (Condition 1, Figure 2) represent any degradation that occurs during the time of sampling and when samples are left outside for pickup. We tested this condition because in many of our remote studies participants collect their samples the day before the pickup is scheduled, with the stabilized blood kept at room temperature overnight. The data show that there was effectively no difference between these samples and the control frozen at -20° immediately after stabilization, suggesting that there is little-to-no degradation of RNA during overnight ambient temperature storage in remote studies. The next set of samples were exposed to varying temperatures following the overnight room temperature incubation for 6 hours (Condition 2, Figure 2). Samples from this condition simulate stabilized blood that sits at a participant’s front door prior to pickup. The mean RIN scores ranged from 6.4 to 7.2 across the different temperatures and only one sample at 25°C yielded a RIN of 4.9 and one at 50°C had a RIN of 5.2. We include 25°C as one of the temperature conditions for incubation to capture study participants who either leave their sample to be picked up indoors in an apartment lounge or mail room or choose to drop their sample off at a UPS store. Finally, condition 3 had an additional overnight room temperature incubation as a representation of specimens that do not get picked up the day following sampling. Samples from this condition replicate situations where the samples are left out for an entire day prior to being returned indoors for another overnight cycle. These samples matched closely to condition 2, further indicating that additional storage at room temperature (up to 38 hours) results in negligible degradation of RNA in homeRNA samples.

**Figure 2.**
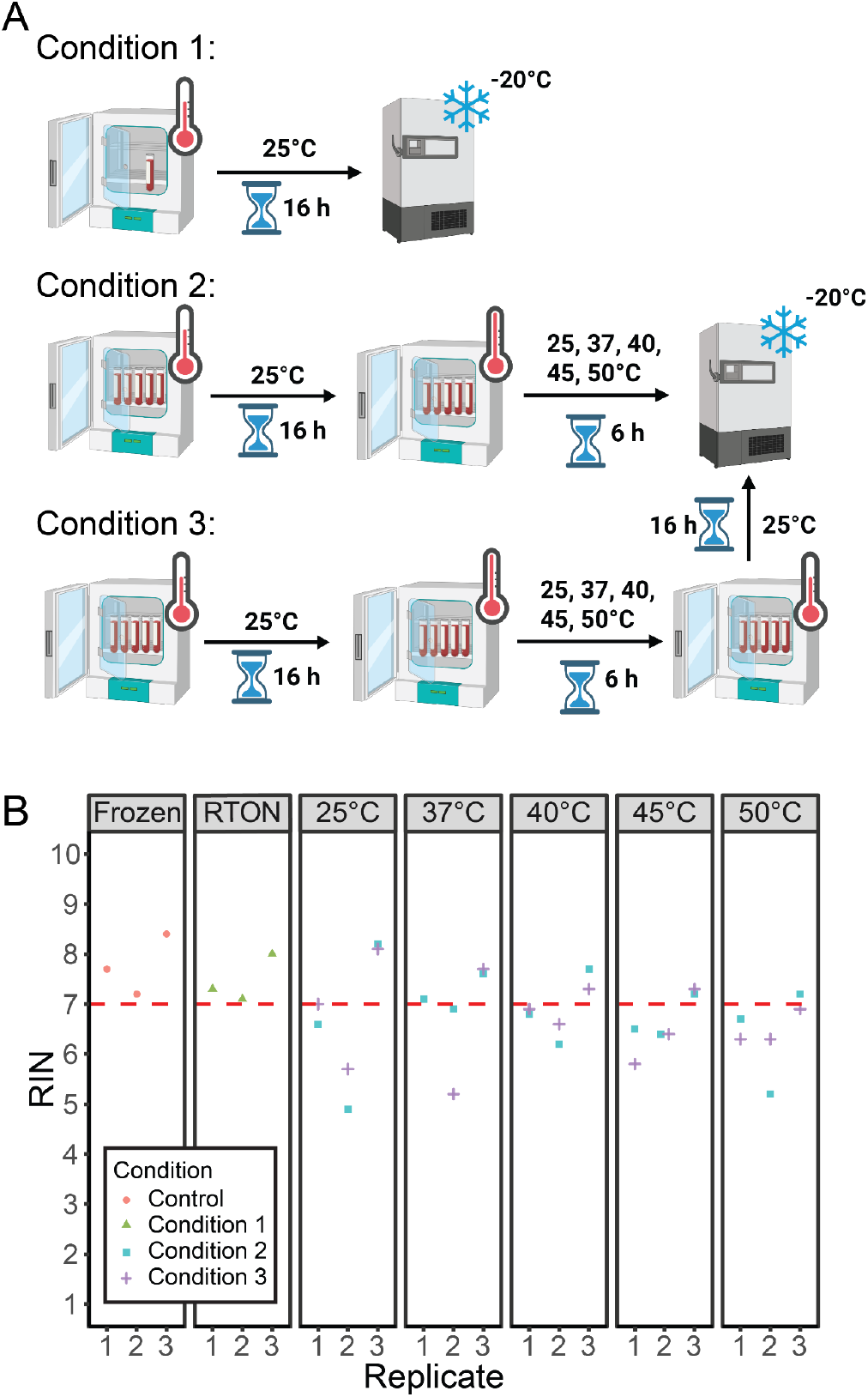
Exposure of RNA*later*-stabilized whole blood samples to shorter term (<2 days) high temperatures (>37°C). (A) Outline of experimental conditions. One sample was frozen at -20°C immediately after stabilizing with RNA*later* to be used as a baseline control. Condition 1 kept one sample at 25°C for 16 hours and then frozen at -20°C; this is referred to as room temperature overnight (RTON). Condition 2 incubated samples at 25°C for 16 hours and then exposed one sample each to 25°C, 37°C, 40°C, 45°C, and 50°C for 6 hours. Condition 3 incubated samples at 25°C for 16 hours and then exposed one sample each to 25°C, 37°C, 40°C, 45°C, and 50°C for 6 hours. After incubation at the different temperatures, all samples were incubated at 25°C for 16 hours then frozen at -20°C. Created with BioRender.com. (B) Resulting RNA quality of each extracted sample from each condition. Each condition was performed in triplicate. Amongst each replicate, all samples across all conditions were extracted in the same batch, resulting in an extraction batch with 12 samples (immediately frozen, one sample from condition 1, 5 samples from condition 2, and 5 samples from condition 3).

### Shipping of RNAlater-Stabilized Blood Across United States with Continuous Temperature Monitoring

While the in-lab temperature-controlled experiments capture a large range of time scales, we were interested in monitoring continuous temperature fluctuations that would be experienced in real shipping conditions (e.g., packages moving from doorstep to shipping vehicle, samples moving between air-conditioned areas to areas without air conditioning, etc.). To this end, we designed an experiment where blood aliquots from a single venous draw were stabilized using the homeRNA kit and shipped out to volunteers across 14 different states. These states are colored green in Figure 3A and include Washington (WA), New Mexico (NM), North Carolina (NC), Minnesota (MN), Maine (ME), Massachusetts (MA), Kansas (KS), Illinois (IL), Georgia (GA), Colorado (CO), California (CA), Arizona (AZ), Hawaii (HI), and Nebraska (NE). Samples were shipped during the summer in August of 2022. We aimed to include a diverse set of states reaching most regions of the U.S. including those located in regions with warm summers (HI, CA, AZ, GA, and NC). The locations include both urban and rural locations, reflected by the Rural-Urban Commuting Area (RUCA) codes (Table 1).^46^ A RUCA value of 1 represents a metropolitan area where a value of 10 represents a rural area. Additionally, our samples were also shipped to an island in Hawaii, which requires longer shipping times to and from Seattle, Washington (location of study lab) than locations within the contiguous U.S.

**Figure 3.**
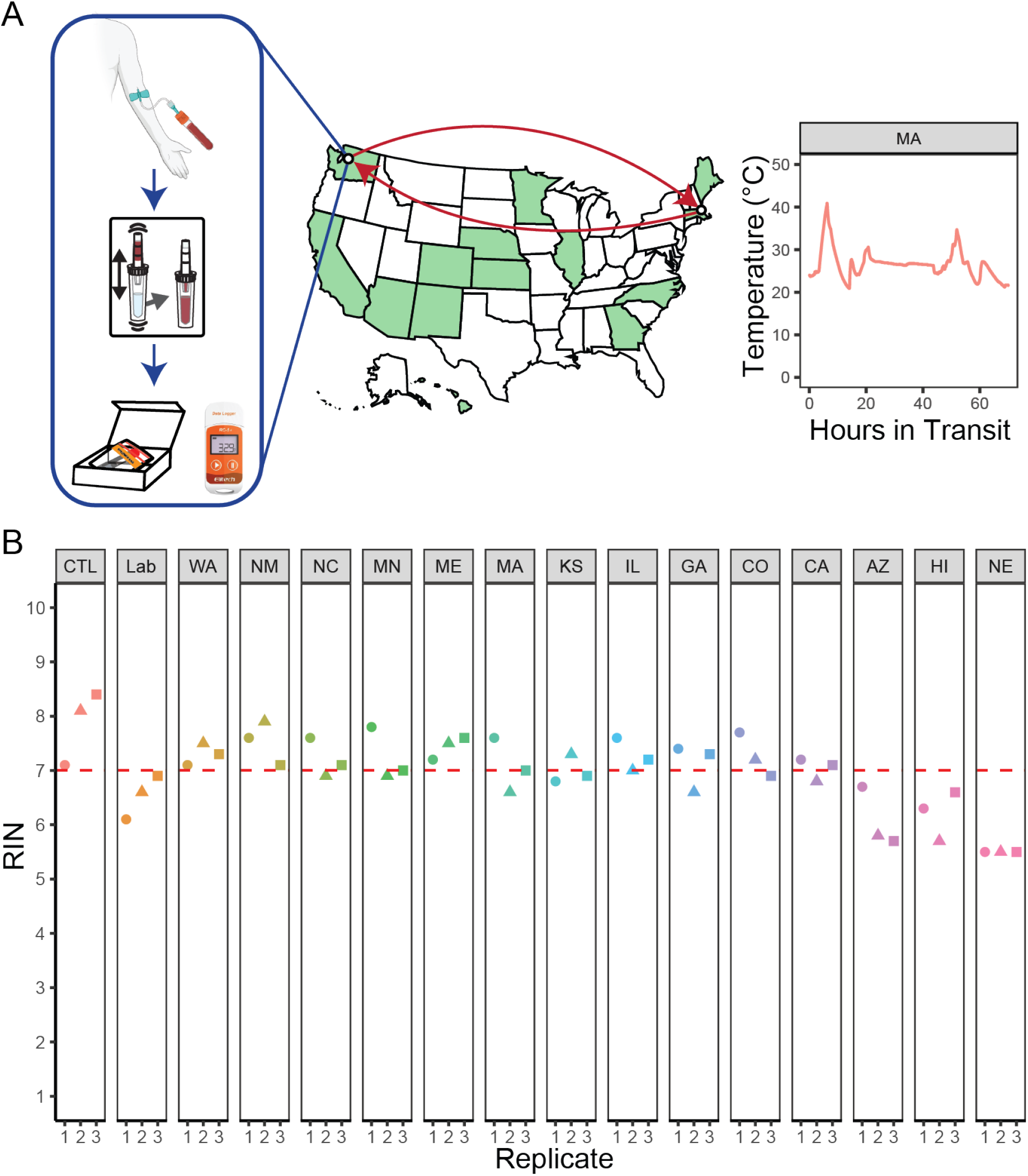
All samples across different shipping locations had RINs suitable for RNA sequencing. (A) Blood from a venous draw from a single participant was aliquoted into the Tasso blood collection tube and stabilized using the homeRNA kit in triplicates and shipped to 14 different U.S. states. The homeRNA kits were packaged as in our other homeRNA studies within a cardboard box and mailer bag, and a continuous temperature probe was directly attached to the samples.^9–11,20^ Once returned to the lab, the RNA was extracted from the blood and tested for integrity. Created with BioRender.com. (B) The resulting RINs are plotted categorized by shipping location. The shape of the data point (circle, triangle, square) corresponds to the extraction batch. Control condition (CTL) was immediately frozen after stabilization, while the in-lab condition (Lab) was kept at 25°C until frozen at -20°C at the time the last sample was received.

Each package also included an Elitech RC-5+ continuous temperature probe (Figure S1) that recorded the temperature from the moment the samples were prepared for shipment in the lab, through the shipping process to the volunteer, to the samples being returned to our lab. These temperature data are plotted in Figure S3 and summarized in Table S1. The data from the temperature probes captures the variable temperature and length of transportation the different samples experienced. The maximum temperature spike we observed was 45.1°C in the sample shipped to NE, which was the most rural of our shipping locations (highest RUCA score) and took 7 days to be returned to the lab. Although HI did not have as high of a RUCA score, the samples shipped to this location took the longest to return to the lab (8 days). Most shipping locations (10 out of the 14 states) took 3 days to return to the lab.

Further, the RIN values from different locations followed a similar trend as the stabilized blood in the temperature-controlled experiments, where longer times and exposure to higher temperatures resulted in lower RIN values. Notably, AZ, NE, and HI had the lowest mean RINs (6.1, 5.5, and 6.2, respectively) and were the locations that took the longest to be returned to the lab. In addition, all three of these states reached temperatures greater than 40°C and spent the longest time above 30°C, with the samples sent to NE spending 55 h above 30°C. Conversely, locations such as CA had similar exposure to a maximum temperature of 44.4°C but had a RIN of 7.0. Even though the maximum temperatures were the same, the longer times in transit and higher median temperatures are the likely causes for the lower RIN from the NE samples.

In the context of remote studies, the data obtained from the real-world shipping experiment is both encouraging and informative. Even the most degraded sample (NE, RIN = 5.5) had RNA of sufficient quality for 3’RNA sequencing (see 3’ mRNA-seq section below). Additionally, all samples were shipped in two directions (to the participant and back to the lab), whereas samples collected in remote studies using homeRNA are only shipped from the participants back to the lab. As such, we expect less degradation to occur in other homeRNA studies since the samples will only be shipped in one direction.

### 3’ mRNA Sequencing of Real-World Shipping Experiment

Finally, we look for selective degradation of RNA fragments from 3’mRNA-seq data. Since all samples in the shipping experiment came from a single blood draw, we can assume that any differences in the measured transcriptome can be attributed to shipping-related degradation. To probe this question further, we selected 20 samples to perform 3’ mRNA-seq analysis. As described in the Experimental section, these included all samples from extraction batch 3 which had the greatest proportion of the median RIN values amongst the three replicates from each location. Additionally, we chose to sequence all three control samples to establish a baseline as well as all three samples from the location with the most degraded samples (NE).

Our first approach is to reference the sequencing data of the samples against a well-characterized immune response signature from the BloodGen3 module repertoire.^28,47^ Each BloodGen3 module consists of a set of genes ranging from 12 to 169 with an average of 37.1 genes per module; these modules have been previously associated with responses to immune-modifying therapies or pathogenic processes.^28^ By calculating enrichment scores (which summarizes the prevalence of a given gene in a module) using the gene set variation analysis (GSVA) method for each of the 382 modules across all samples, we aimed to break down the results and provide a more granular understanding of the immune response landscape.^29^ Principal component analysis (PCA) of these scores revealed minimal clustering patterns based on shipping location (Figure 4A), indicating a lack of significant variability within the shipped samples. While the In-Lab samples (kept at 25C for the duration of the experiment, 8 days) did exhibit some clustering, the Control sample (immediately frozen) clustered with the rest of the shipping locations. However, it is important to note that the overall variability captured by the PC1 is only 16.25% and PC2 is 8.97% and that the distribution of the points is relatively tight. The proximity of the Control to the rest of the samples also indicates that there is very little variability arising from the shipping process. The heatmap visualization of enrichment scores across various modules and samples (Figure 4B) and the focused analysis of three key module groups (module aggregates A28, A35, and A37, corresponding to interferon, inflammation and circulating erythroid cells signatures, respectively) (Figure 4C) further supported this observation, with only minor variations in enrichment scores across individual samples. These findings highlight the relative homogeneity of our samples and indicate little variation depending on the sample location.

**Figure 4.**
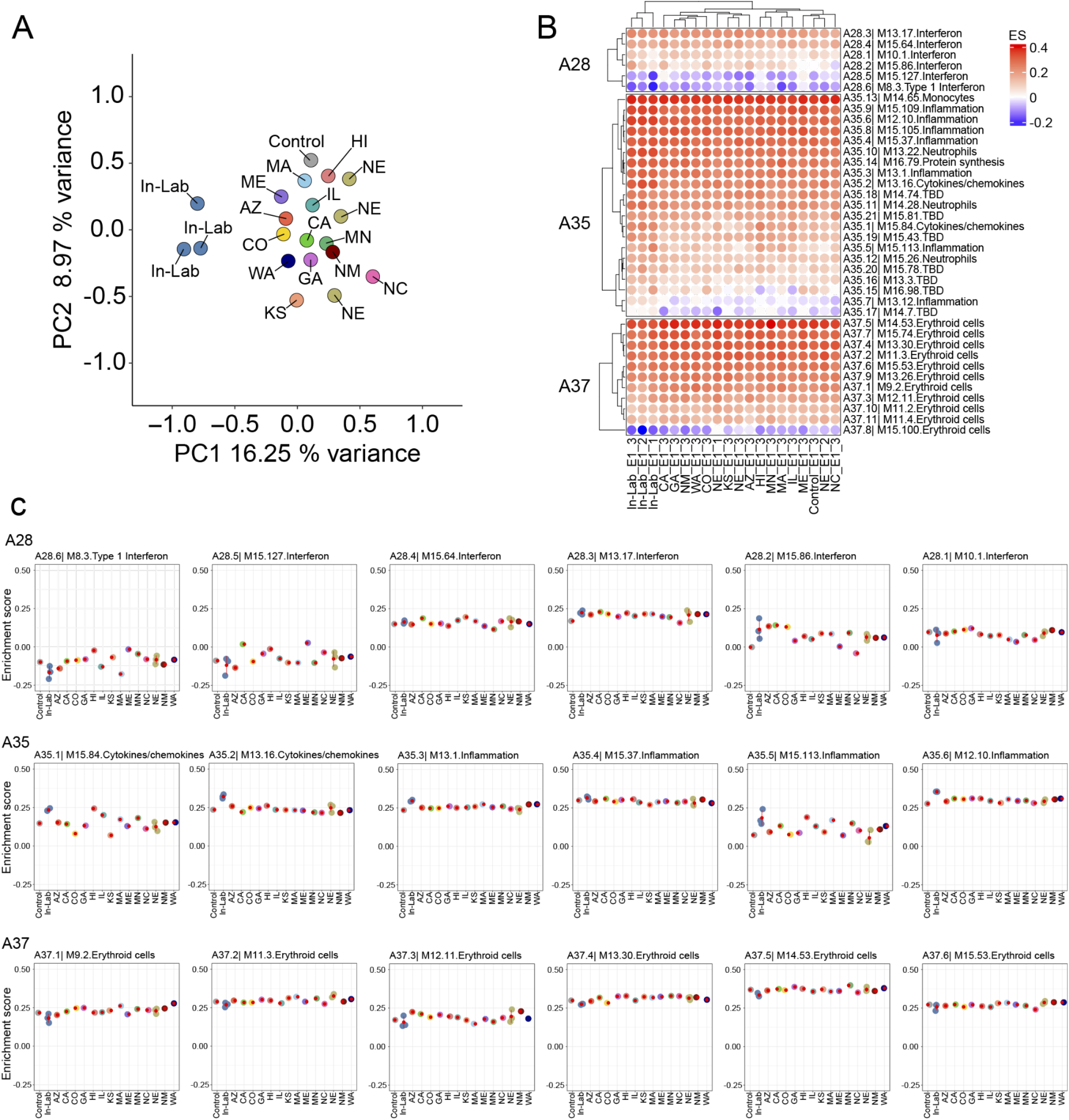
Analysis of Module Enrichment Scores in Samples. (A) PCA of sample data by location. This PCA plot provides a visualization of the data variance, with each point representing a sample’s projection on the two principal components labeled based on sample condition (shipping location, control (frozen immediately), or in-lab sample (stored at 25°C for 8 days)). (B) Heatmap of enrichment scores across conditions. Enrichment scores for various biological modules are depicted in a heatmap format, with rows representing different modules (e.g., interferon response, inflammation) and columns representing individual samples. The color gradient indicates the level of enrichment from decreased (blue) to increased (red). (C) Detailed module enrichment scores for groups A28, A35, and A37. The dot plots show the enrichment scores for selected modules within the three groups. Individual sample scores are plotted on the x-axis against their corresponding enrichment score on the y-axis.

Further, to contextualize the observed variation in gene expression in our samples, we include results from a previously published COVAX study, which investigated the immune response to COVID-19 mRNA vaccine.^30^ By comparing the enrichment scores of six interferon response modules (M8.3, M10.1, M13.17, M15.64, M15.86, and M15.127, comprised in the module aggregate A28) at different timepoints before and after vaccination, we can observe the differences in signal variability occurring due to a well-controlled biological signal. The dot plot in the supporting information (Figure S4) demonstrates a clear increase in interferon module enrichment scores following vaccination, with peak responses observed at days 2 and 3. This distinct pattern of vaccine-induced immune activation serves as a reference point for interpreting the variability observed in our samples (labeled as UW in Figure S4). Assuming that the stabilized blood exhibits minimal DNA transcription *ex vivo*, any large difference in transcript counts can be attributed to degradation during the shipping process. The tight clustering of our samples compared to the large differences in the COVAX data suggests that the variability introduced by temperature exposure during shipping is much smaller in magnitude than the changes induced by a biological signal, such as an immune response elicited by vaccination. Taken together, this indicates that there is minimal RNA degradation of homeRNA-stabilized samples owing to variable exposure to temperature and shipping times.

We also fit a model that includes the time spent above a temperature of 30°C, RIN, and 4 surrogate variables (generated using the Bioconductor sva package) to account for any unobserved technical variability.^34^ After fitting this model, we selected genes with a false discovery rate (FDR) of <0.05, which estimates the maximum number of false positives in a set of significant results, as well as requiring at least a 1% change in expression. This resulted in only two genes (MMP9 and ADGRG3) that had a decrease in expression as a function of time spent at 30°C or higher (Figure S5). While both genes appear to be affected by storage at high temperatures, they also exhibit a notable amount of noise mainly due to the small sample size. Moreover, linear regression is not resistant to outliers; in our data, we propose that there are some (3 for MMP9 and 4 for ADGRG3) high-leverage points that are multiple orders of magnitude greater than the rest of the points and are not representative of the rest of the samples. This suggests that the two genes MMP9 and ADGRG3 potentially could be affected by shipping conditions, but future work with more sensitive methods could potentially capture differences in the transcriptome not detected here.

Further, we fit linear regressions with “time in transit” or “time above 30°C” against RIN scores to better understand the effects these variables have on RNA degradation (Figures S6 and S7). All samples that spent below 100 hours in transit had RIN scores at or above 7. The samples that were in transit the longest were AZ (168 h), HI (193 h), and NE (168 h) and had mean RINs of 6.1, 6.2, and 5.5. Here, the data show that the time in transit alone is not representative of RIN since HI had a higher RIN score than NE even though it was in transit for longer. If we consider time spent over 30°C, AZ was exposed to temperatures above 30°C for only 38.3 hours, HI was exposed for 22.8 hours, while NE was exposed for a total of 55 hours. When the time spent above 30°C is considered, the corresponding RIN scores seem to follow a more sensible pattern with longer exposures leading to lower RIN values.

In summary, based on our 3’ mRNA-seq dataset, there is little evidence that differences in shipping conditions result in consistent changes in the measured mRNA expression levels in blood RNA from a single donor. We expect that samples in other remote studies using homeRNA that have similar RINs will also exhibit negligible transcript-specific degradation from variable shipping conditions as measured by 3’ mRNA-seq. It is important to note that traditional RNA-seq methods may yield different results owing to differences in read depth and genes detected. Further work is needed to conclusively determine if the homeRNA system is completely resilient to preferential degradation of transcripts when exposed to variable shipping conditions, however, the current work shows promising results.

## Conclusion

We expand upon existing literature on RNA stability in cold storage conditions and dried blood spots by investigating the effects of variable temperature exposure on RNA*later*-stabilized samples, with a systematic study on the effects of high temperature exposure and shipping time on liquid blood samples. In the temperature-controlled experiments, we found that longer storage times (4 and 8 days) at high temperatures (>37°C) resulted in lower RIN values, while shorter exposures (∼6 hours) to high temperatures resulted in minimal RNA degradation. We also used homeRNA-stabilized blood samples in a real-world shipping experiment. All shipped samples were sequenced using Lexogen’s QuantSeq 3’ mRNA technology, with minimal differences in the transcriptome measured.

This study illustrates the utility of the homeRNA platform in high temperature settings. It also serves as a guide for other researchers who aim to conduct remote sampling studies that use RNA as a readout and rely on shipping via courier service for sample retrieval. Our findings strongly suggest that there is limited preferential degradation of transcripts in the shipping process and, as such, give us confidence in interpreting transcriptomic data from homeRNA-stabilized whole blood samples. These results will inform ongoing homeRNA studies and will be critical for future study design. Future work will use larger sample sizes, longer read depth sequencing such as total RNA-seq to further substantiate these findings, and work in low- and middle-income countries (LMICs) with different courier methods and climates.

## Acknowledgements

This publication was supported by the National Institutes of Health (NIH) through the University of Washington EDGE Center of the National Institute of Health (P30ES007033), R21ES034338, and R35GM128648. The content is solely the responsibility of the authors and does not necessarily represent the official views of the National Institutes of Health. We would like to acknowledge Fang Yun Lim for the helpful conversations regarding experimental design, the logistics of our real-world experiment, as well as the many contributions to prior work that helped inform the current study. We would also like to acknowledge the 14 volunteers who helped us with our real-world shipping experiment and the study participants who participated in venous blood draws.

