## Supplemental Information for "Your Blood is Out for Delivery: Considerations of Shipping Time and Temperature on Degradation of RNA from Stabilized Whole Blood" (from another version of this preprint: version 1, 2024-08-25)

### EXPERIMENTAL

#### Participant Recruitment and Demographics

Healthy adult volunteers (18 years or older) were recruited in Seattle, Washington via word of mouth under a protocol approved by the University of Washington Institutional Review Board study number STUDY00014133. Written informed consent was obtained from all participants.

#### RNA Isolation

For all samples, total cellular RNA was isolated using the Ribopure Blood RNA Isolation Kit (Thermo Fisher) according to manufacturer's protocol and eluted in two 50 µl aliquots. Isolated RNA was stored at -80°C until ready for further analysis.

#### Investigating RNA Integrity of RNA*later* Stabilized Blood Samples in Temperature Controlled Experiments

*Effects of a Freeze-Thaw Cycle on RNA Stability*

For this experiment, blood from a venous blood draw is stabilized using RNA*later* according to manufacturer instructions. The blood is then aliquoted into 9 samples which are separated into three conditions: (1) control which is immediately frozen, (2) room temperature (25°C) for 24 h (room temperature overnight, RTON) then frozen, and (3) freeze-thaw (FT) where the samples were frozen immediately after stabilization for 24 h, then kept at room temperature for 24 h before being frozen again. The samples were frozen at -20°C for the duration of the experiment then moved to -80°C until further processing. The experiment was performed two more times for a total of three replicates. Once the RNA was extracted the samples were tested for RNA integrity using the Agilent 2100 Bioanalyzer.

#### Assessment of total cellular RNA integrity and yield

After isolating the RNA from the homeRNA-stabilized samples, RNA Integrity number (RIN) values were obtained on a Bioanalyzer 2100 (Agilent) from the first 50 µl elution. The Agilent 2100 Bioanalyzer uses proprietary, built-in software to calculate RIN values, which is based on the relative peak height and shape of the 18S and 28S rRNA fragments in the resulting electropherogram after separation. All samples were assessed using the RNA 6000 Nano Kit (Agilent).

For the in-lab temperature controlled experiments, RNA concentration was measured from the first 50 µl elution aliquot using a Cytation 5 Multi-Mode Reader (Agilent Biotek Instruments) with a Take3 Micro-Volume Plate at the wavelengths of 260, 280, and 320 nm. For the field experiments, RNA concentration was measured from the first 50 µl elution using a Qubit 4 Fluorometer using the RNA High Sensitivity (HS) Assay kit (Invitrogen).

**Investigating RNA Integrity of RNA*later* Stabilized Blood Samples in Real World Setting with homeRNA**

*Visualization of Temperature Over Time Across Shipping Locations*

The data from the temperature probe directly attached to the samples is assumed to have readings closest to the "true" temperature the samples would experience in transit and only the data from this probe are visualized and used for downstream analysis. Since the temperature probes were not stopped until all packages arrived, the data needed to be filtered prior to graphing. The data were filtered using the R dplyr package such that the time did not exceed 200 hours (close to the maximum time for the last sample returned to lab) and was above 10°C given that the lowest temperature reading prior to being frozen was closer to 20°C for all samples. The temperature was then plotted against time (hours) to illustrate the variability in shipping process and temperature fluctuations.

**TABLES**

**Tables 1: Summary of temperature probe data**

| Location | RUCA | Time (h) | $T_{avg}$ (℃) | $T_{med}$ (℃) | $T_{max}$ (℃) | $T_{sd}$ (℃) | Hours above 30℃ |
|---|---|---|---|---|---|---|---|
| AZ | 1 | 168 | 26.8 | 26.9 | 40 | 4.2 | 38.3 |
| CA | 1 | 72 | 26.5 | 25.4 | 44.4 | 4.9 | 11.1 |
| CO | 4 | 96 | 24.3 | 24.1 | 39.5 | 4.4 | 7.6 |
| GA | 1 | 72 | 24.8 | 24.3 | 41.9 | 4.4 | 7.4 |
| HI | 4 | 193 | 26.1 | 25.5 | 38.5 | 3.7 | 22.8 |
| IL | 1 | 72 | 23.7 | 22.6 | 38 | 1.9 | 4.5 |
| In-Lab | 1 | 200 | 25.5 | 25.5 | 25.7 | 0.4 | 0 |
| KS | 1 | 71 | 25.8 | 24.0 | 43.3 | 4.9 | 10.4 |
| MA | 1 | 72 | 26.4 | 26.5 | 40.9 | 3.5 | 8.2 |
| ME | 5 | 72 | 24.0 | 23.4 | 35.4 | 3.2 | 3.8 |
| MN | 1 | 71 | 25.4 | 23.3 | 39.3 | 5.6 | 14.4 |
| NC | 5 | 72 | 27.7 | 28 | 38.8 | 4.1 | 18 |
| NE | 10 | 168 | 28.5 | 27.2 | 45.1 | 5.5 | 55 |
| NM | 1 | 72 | 26.4 | 26.1 | 39.3 | 3.9 | 9.5 |
| WA | 1 | 71 | 23.8 | 23.1 | 36.6 | 3.1 | 3.7 |

### FIGURES

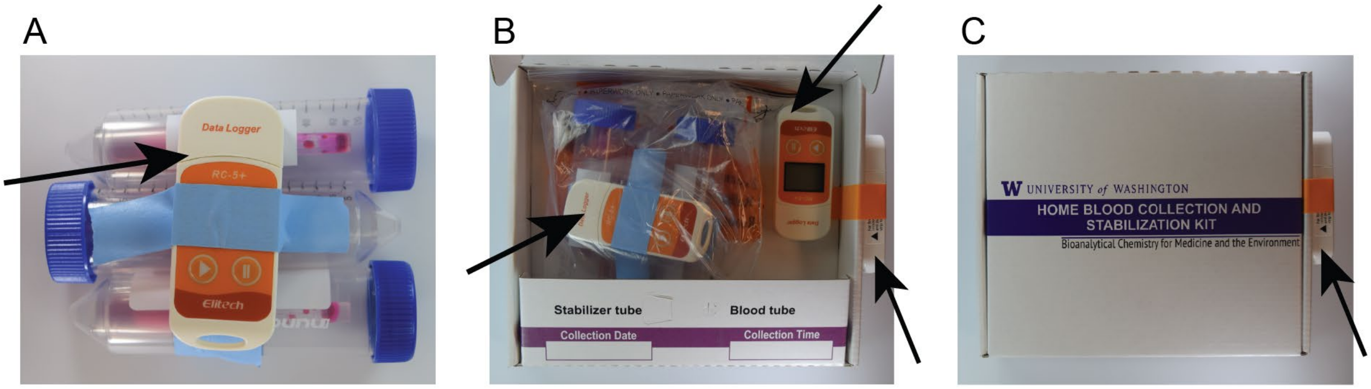


**Figure S1**. Shipping experiment continuous temperature probe setup. (A) Each homeRNA-stabilized sample is placed in a 50 mL conical tube with a custom insert to prevent excessive movement in the shipping process. Three such replicates are taped together and a temperature monitor (indicated by black arrows) is attached directly to this construct. (B) The samples are sealed in a biohazard bag and are then placed in the homeRNA box. Another temperature probe is placed inside of the box as a backup. (C) Then an additional temperature probe is attached to the outside as a final backup; the entire box is placed into a UPS LabPak and shipped to the participants. These samples are kept indoors overnight at which point a pre-printed return label (in a label pouch on the outside of the LabPak) is attached. The LabPak is placed on the volunteers' front porch or mailroom and picked up by UPS to be returned to the lab.

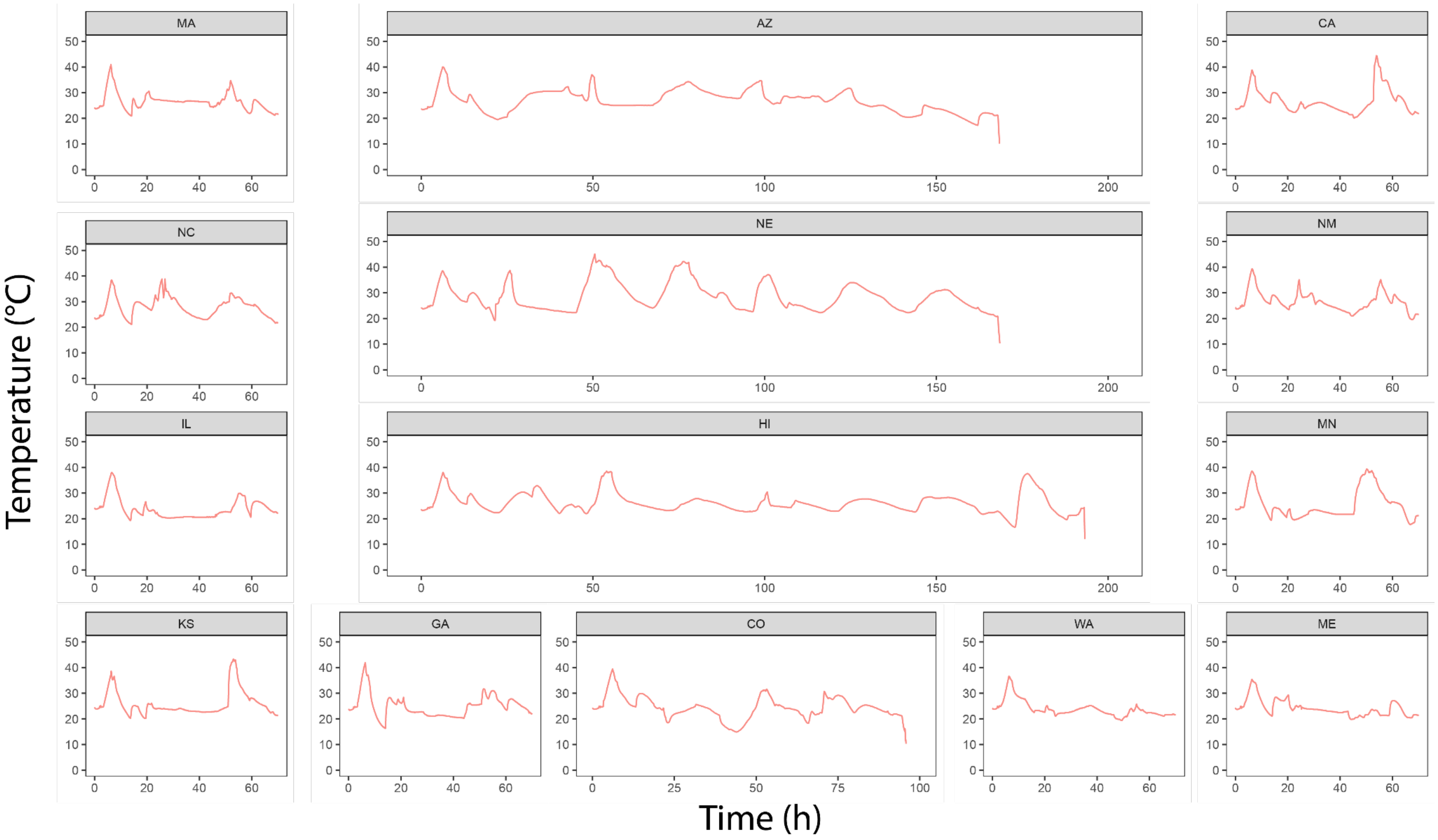


**Figure S2**. Summary of temperature probe data from each location.

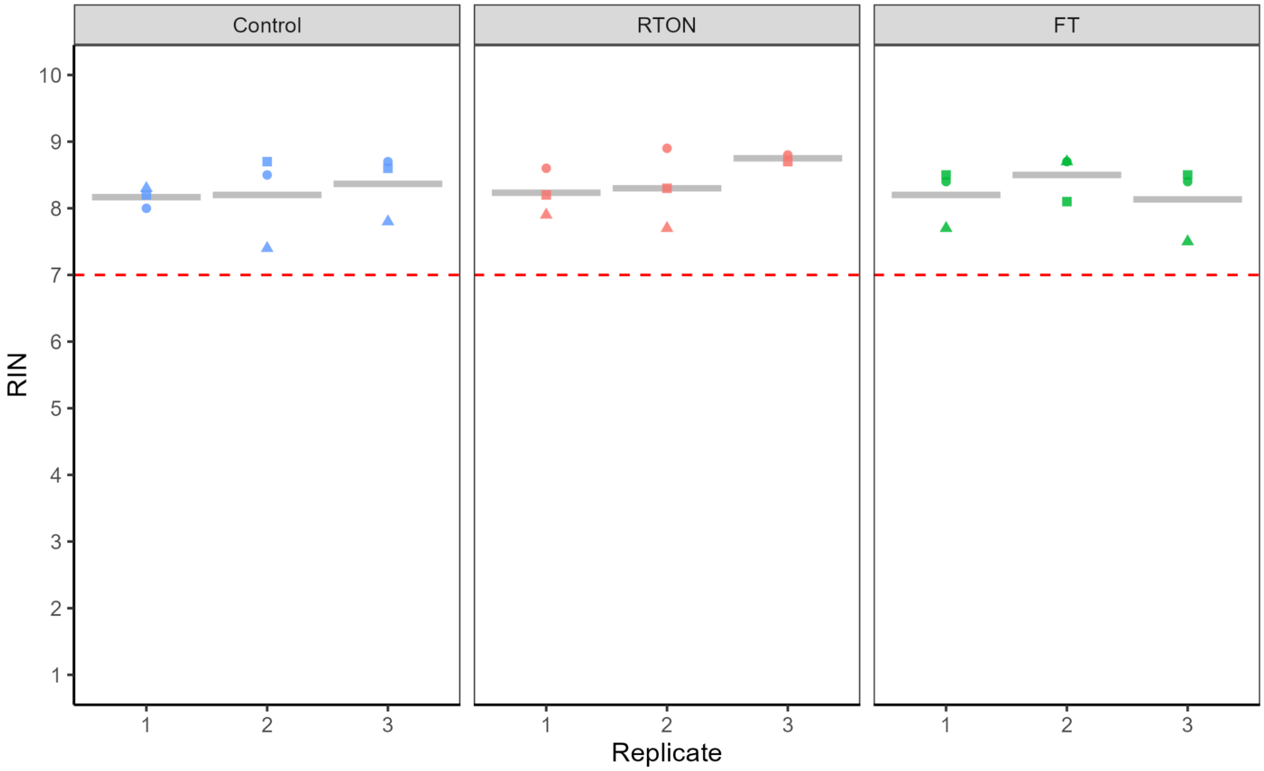


**Figure S3**. A single Freeze-Thaw cycle has negligible effects on RNA integrity in homeRNA stabilized samples. The resulting RINs are plotted here where each column represents the condition, with the gray crossbars indicating average values for each replicate. Overall, the data did not suggest that there were any adverse effects on RNA integrity as a result of a freeze-thaw cycle of homeRNA-stabilized blood samples.

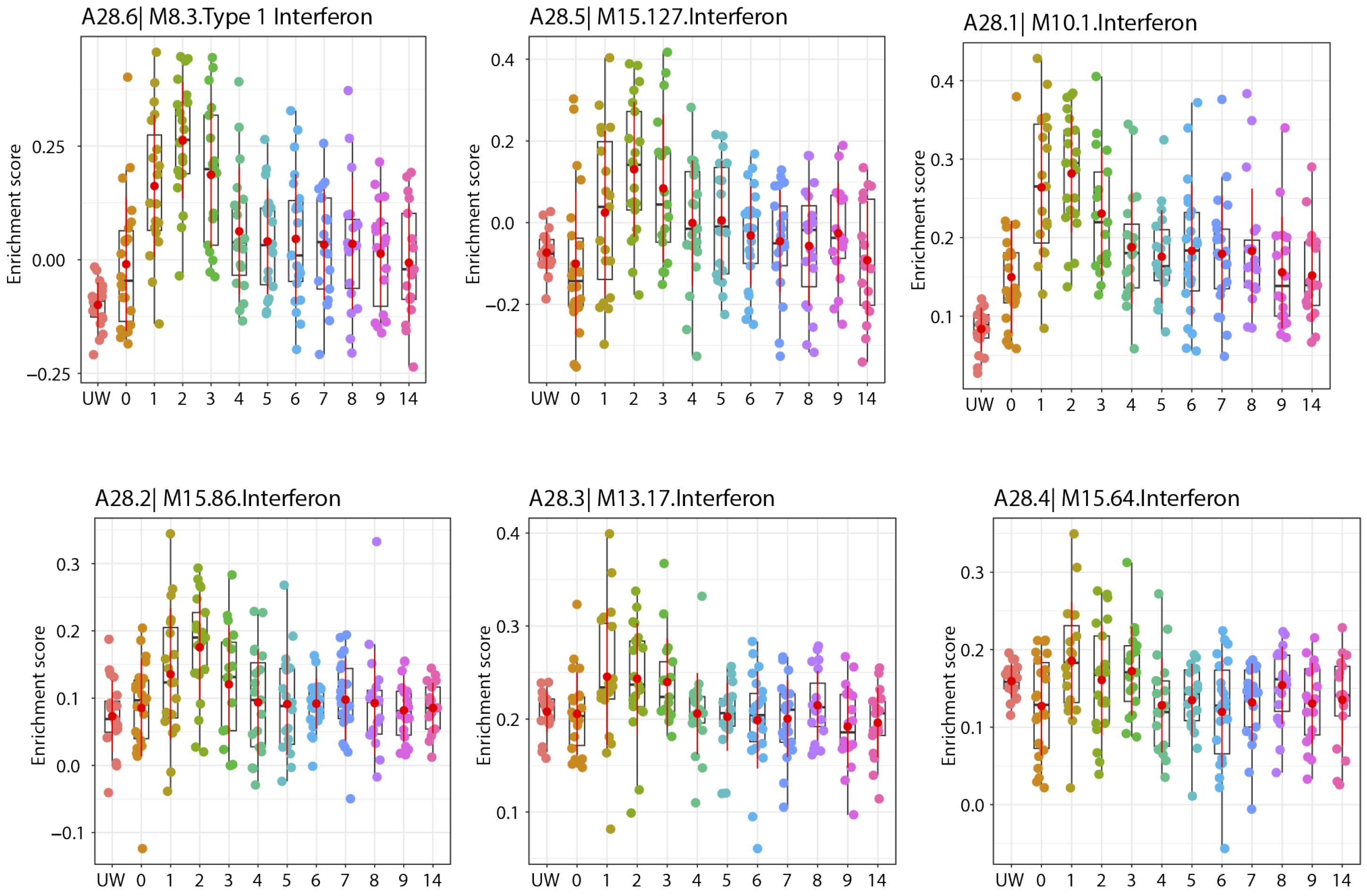


**Figure S4**: Analysis of Interferon Modules Enrichment Scores of Subjects Pre- and Post- Vaccination. The dot plots show the enrichment scores for six interferon response modules (M8.3, M10.1, M13.17, M15.64, M15.86, and M15.127) for the current study (UW) and at different days before and after administration of COVID-19 mRNA vaccine (COVAX study – day 0 = pre-vaccination sample). Following the UW (shipping) samples, the y-axis represents days of sampling from the COVAX study.

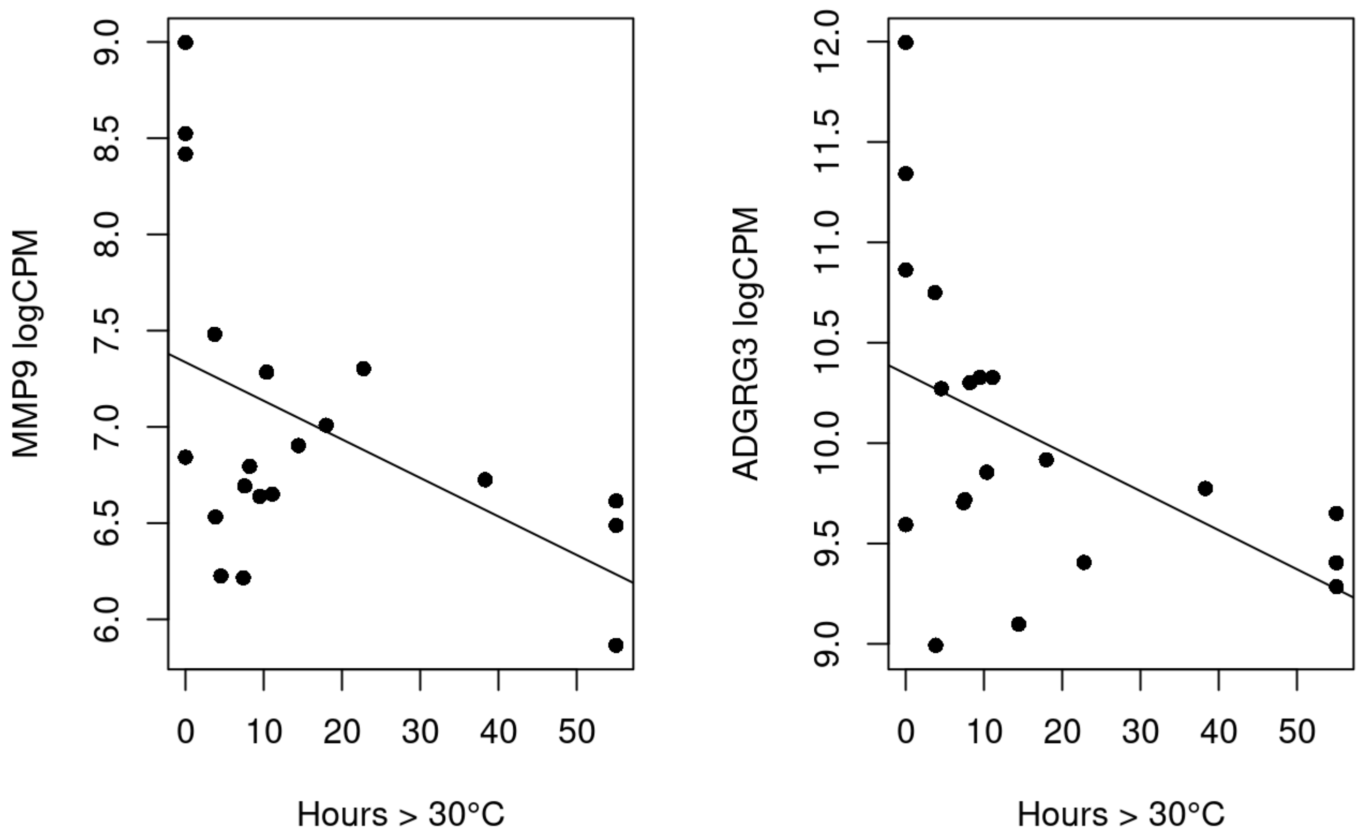


**Figure S5**. Expression levels of MMP9 and ADGRG3 as a function of time in excess of 30°C. The slope for both genes corresponds to an approximate 34% reduction in transcript for each 10 hours > 30°C.

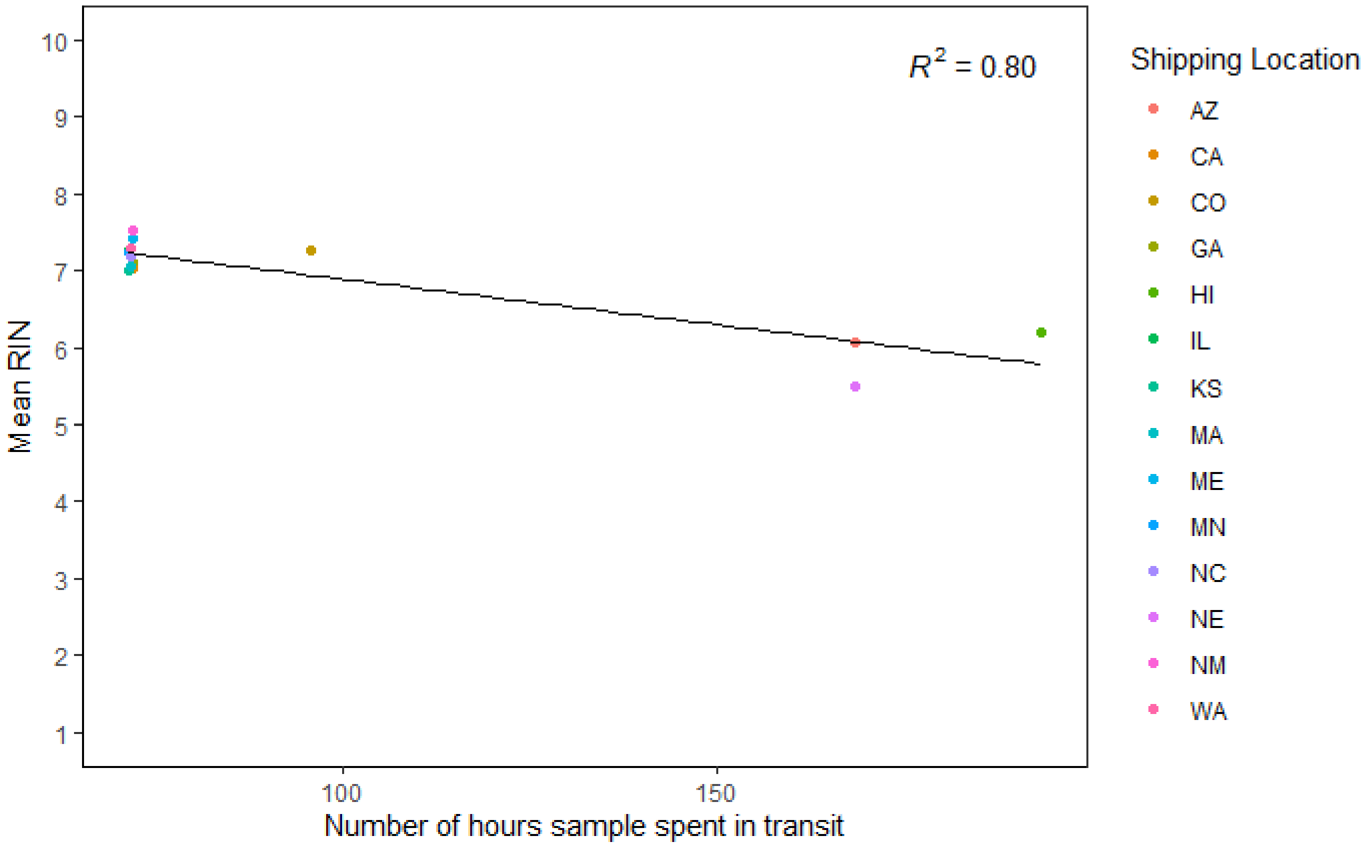


**Figure S6**. Increasing time in transit minimally affects the RNA integrity of homeRNA-stabilized blood samples. Mean RIN scores for each location are plotted against the times in transit for each location which are extrapolated from the temperature probe data visualized above. Though a trend of lower RINs with longer times in transit is observed, the $R^2$ value is only 0.57, indicating that there is not a strong correlation between the two variables. Despite this trend, all RIN values are well within the acceptable range for 3' mRNA sequencing.

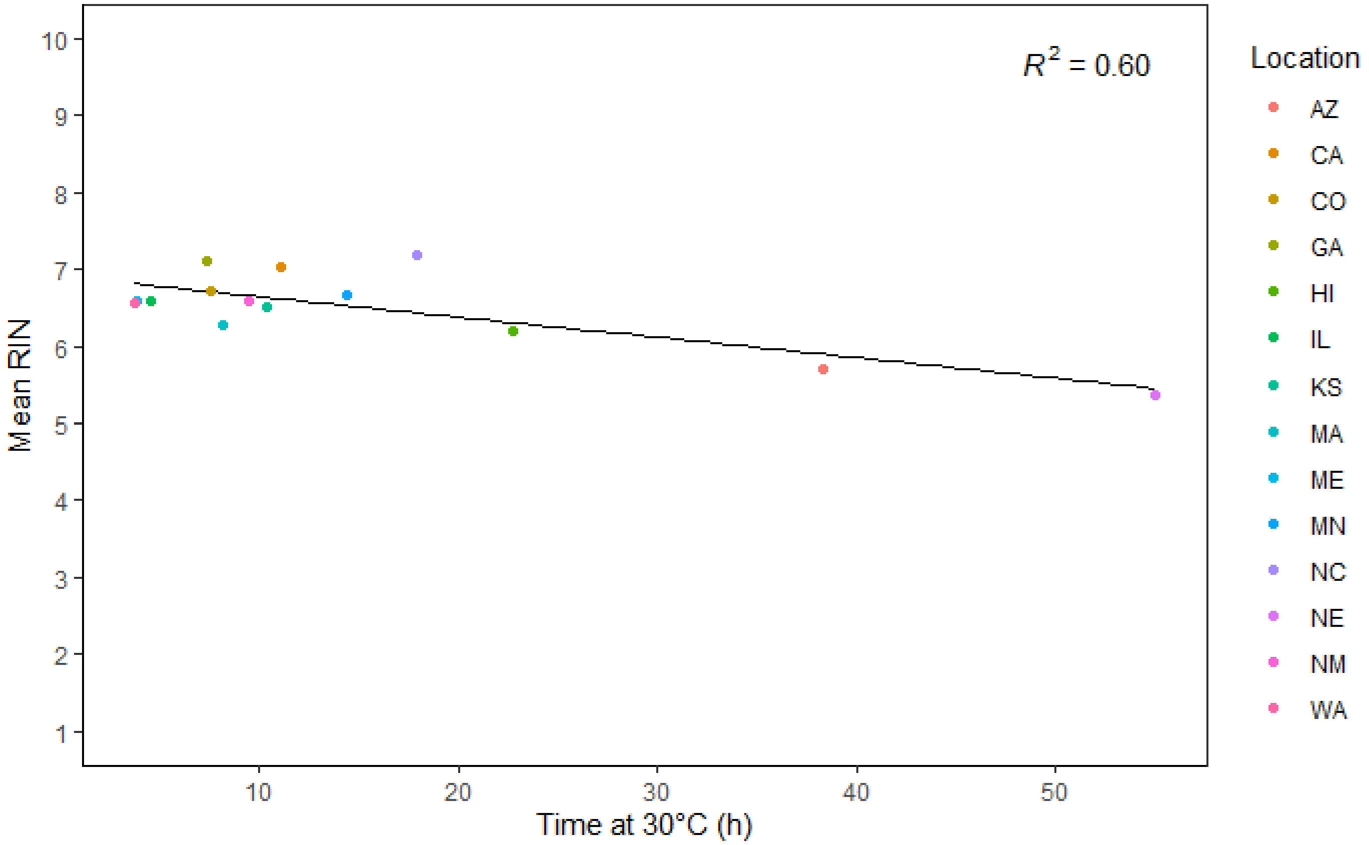


**Figure S7**. Longer times spent above 30°C minimally affect the RNA integrity of homeRNA-stabilized blood samples. Mean RIN scores for each location are plotted against the number of hours a sample experienced a temperature of 30°C or higher. Longer times spent at or above 30°C correspond to lower mean RIN values, however, there is not a strong correlation between the two variables ($R^2$ = 0.60)

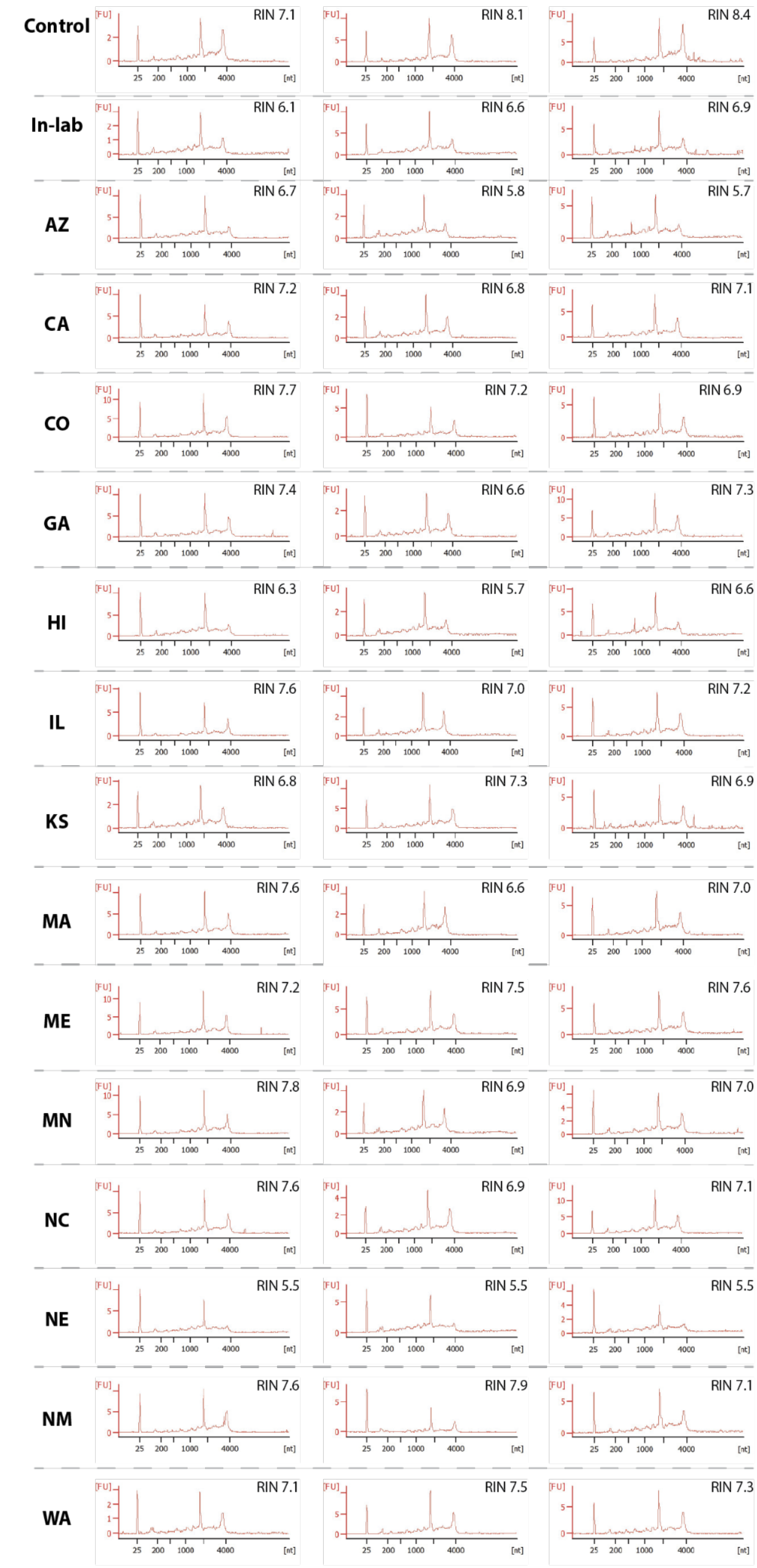


**Figure S8**. Electropherograms of isolated RNA from RNA*later*-stabilized blood samples shipped to 14 different US states. Electropherograms were obtained as raw data using a RNA 6000 Nano Kit on an Agilent 2100 bioanalyzer, which uses fluorescence to detect the marker (~25 nt), 18S rRNA fragments (~1900 nt), and 28S rRNA fragments (~3900 nt). The RIN algorithm then uses these peaks to assign a RIN value to the corresponding sample. Each state was shipped blood samples in triplicate (left, middle, and right columns are replicates 1, 2, and 3, respectively). Samples from the third replicate (right column) were sequenced with 3'mRNA-seq. All three replicates from the Control and Nebraska (NE) samples were also sequenced with 3'mRNA-seq.
